# Progressive spreading of DNA methylation in the GSTP1 promoter CpG island across transitions from precursors to invasive prostate cancer

**DOI:** 10.1101/2022.12.21.521517

**Authors:** Harshath Gupta, Hitoshi Inoue, Yasutomo Nakai, Masashi Nakayama, Tracy Jones, Jessica Hicks, Balasubramanian Kumar, Meltem Gurel, William G. Nelson, Angelo M. De Marzo, Srinivasan Yegnasubramanian

**Author notes:** Corresponding authors: AMD,; SY. These authors contributed equally (Harshath Gupta, Hitoshi Inoue).

## Abstract

Glutathione S-transferase pi 1 (GSTP1) is lowly expressed in normal prostate luminal cells and becomes induced in most proliferative inflammatory atrophy lesions (PIA). GSTP1 becomes silenced in prostatic intraepithelial neoplasia (PIN) and prostate adenocarcinoma (CaP) via cytosine-phospho-guanine (CpG) island promoter hypermethylation. However, *GSTP1* methylation patterns in PIA and PIN, and their relationship to patterns in CaP are poorly understood. We used bisulfite genomic sequencing to examine patterns of *GSTP1* promoter CpG island methylation in laser capture microdissected benign, PIA, PIN, and CaP regions from 32 subjects that underwent radical prostatectomy. We analyzed 908 sequence clones across 24 normal epithelium, 37 PIA, 18 PIN, and 23 CaP regions, allowing assessment of 34,863 CpG sites with allelic phasing. Normal and PIA lesions were mostly unmethylated with 0.52 and 1.3% of total CpG sites methylated, respectively. PIN and CaP lesions had greater methylation with 24% and 51% of total CpG sites methylated, respectively. The degree of *GSTP1* methylation showed progression from PIA ≪ PIN < CaP. PIN lesions showed more partial methylation compared to CaP lesions. Partially methylated lesions were enriched for methylation changes at AP1 and SP1 transcription factor binding sites. These results demonstrate that methylation density in the *GSTP1* CpG island in PIN was intermediate relative to that in normal prostate epithelium/PIA and CaP lesions. These results are consistent with gradual spreading of DNA methylation centered at the SP1/AP1 transcription factor binding sites in precursor lesions, with subsequent spreading of methylation across the entire CpG island in transition to CaP.

## Background

Initiation of cancer involves multiple genetic events characterized by chromosomal translocations, deletions, amplifications and point mutations of critical genes (1). In addition to these genetic changes, abnormal epigenetic alterations, including changes in patterns of DNA methylation, have been observed in a wide spectrum of human cancers, including widespread genomic hypomethylation and simultaneous regional increases in DNA methylation (2,3). This aberrant somatic DNA methylation of CpG dinucleotides within upstream regulatory regions of genes is generally associated with a compacted chromatin structure, and accompanying repression of transcription (4). A promising area of strength for DNA methylation in the clinic is in the area of molecular diagnostics and early detection (5). The hypermethylation of promoter CpG islands of tumor-suppressor genes is a common alteration in cancer cells, and leads to the transcriptional inactivation and loss of normal cellular functions (6). In many cases hypermethylation of the CpG island in gene regulatory regions has been correlated with a loss of gene expression; thus it is likely that DNA methylation provides an alternate pathway to gene deletion or mutation for the loss of tumor suppressor gene function.

A large body of evidence suggests that the major precursor lesion to invasive prostatic adenocarcinoma (CaP) is high-grade prostatic intraepithelial neoplasia (HGPIN) (7–10). This hypothesis is supported by findings of proximity to CaP, morphological changes and molecular alterations shared between the two lesions, and two-fold increased prevalence and extent of HGPIN in prostates with carcinoma vs those without. However, recent evidence suggests that at times, lesions morphologically identifiable as HGPIN may also arise as retrograde spread from invasive carcinomas that invade otherwise benign glands and mimic the morphology of HGPIN (11,12). We refer to such lesions as post-invasive intraepithelial carcinomas, or PIC. This is analogous to intraductal carcinoma of the prostate, which represents an aggressive form of prostate cancer that is presumed to arise through such retrograde invasion of invasive cancer into preexisting benign glandular structures. Therefore, it is important to develop methods to determine whether a given lesion with the appearance morphologically of high grade PIN, represents *de novo* high grade PIN or represents PIC mimicking PIN. DNA methylation changes in cancer cells offer the potential to distinguish whether HGPIN lesions represent CaP precursors or CaP itself (e.g. PIC), because somatic CpG dinucleotide methylation patterns at specific genome sites can be sparse or dense, allowing the stepwise molecular progression from initially unmethylated (normal prostate) to ultimately extensively methylated (CaP) alleles, transiting through intermediate methylation in precursor lesions.

The CpG island promoter region spanning the glutathione S-transferase (*GSTP1*) gene is methylated in approximately 90% of prostatic adenocarcinomas. *GSTP1* encodes an enzyme involved in detoxification and protection from oxidants and carcinogens by conjugation to glutathione (13). GSTP1 is expressed at high levels in normal basal cells and is expressed at very low levels in normal luminal cells. However, it appears highly induced in proliferative inflammatory atrophy (PIA), a lesion associated with repeated injury and inflammation, in which many cells express high levels of GSTP1, apparently in response to increased chemical stress. In contrast, HGPIN and CaP are often devoid of GSTP1 protein, and often harbor methylation of the CpG island at the *GSTP1* promoter (4,10,14–16).

As hypermethylation in the 5′ region and promoter of *GSTP1* appears to be a frequent and early event in prostate cancer development, we wished to examine the specific changes in the pattern of methylation spanning the gene promoter in CaP and candidate precursor lesions, HGPIN and PIA. We used bisulfite genomic sequencing to determine the methylation status of 39 cytosine residues present spanning the *GSTP1* gene promoter. Specifically we wished to address whether the hypermethylation (1) proceeded via intermediate molecular steps, (2) was enriched at specific regions in the promoter, such as transcription factor binding sites, and (3) presented with inter- or intra-tumor heterogeneity across lesions within patients.

## Methods

### Prostate Tissue Samples

Frozen or formalin-fixed paraffin-embedded (FFPE) radical prostatectomy specimens were selected from 32 patients undergoing radical prostatectomy at The Johns Hopkins Hospital in which PIA and PIN lesions were readily discernible histologically using H&E staining and encompassed regions large enough to be amenable to laser capture microdissection. No subjects were excluded from our study, and male subjects were eligible for the study if they were undergoing radical prostatectomy for non-metastatic prostate cancer. Also, to avoid the potential that PIN lesions represented intraductal/intra-acinar spread of carcinoma, most of them were isolated from tissue blocks either not containing carcinoma or in which the PIN lesions were “away” from carcinoma by at least 2 mm. Also, since HGPIN can be difficult to diagnose on frozen sections, most of the PIN lesions analyzed were from FFPE sections (12/18). The PIN lesions that we did use from frozen tissues were clearly diagnosable as high grade PIN on those sections. All patients provided informed consent for use of tissues, and the use of tissues was approved by the Johns Hopkins University School of Medicine Institutional Review Board.

### Laser Capture Microdissection (LCM) and Isolation of Genomic DNA

Three serial 6-μm sections were stained with hematoxylin. Matched regions of normal epithelium (4 of the regions of normal were from transition zone in histological BPH nodules), PIA, HGPIN, and CaP were obtained by LCM using the Veritas system (Arcturus Engineering Inc., Mountain View, CA) and the CapSure HS LCM Caps (Arcturus Engineering Inc.). Approximately 1000 to 3000 cells were obtained in most cases by LCM. DNA was isolated with a standard phenol/chloroform protocol. DNA quantification was carried out by real-time PCR of the Beta-globin gene to ensure that DNA was of ample quality for reliable quantitative PCR amplification (15). Some of the sections were stained with antibodies against GSTP1 for immunohistochemistry.

### Bisulfite Genomic Sequencing

Two nanograms of genomic DNA was bisulfite converted using the EZ DNA methylation kit (Zymo Research, Orange, CA) and eluted in 11 μl of TE buffer, pH 7.4. Portions (1 μl) of the DNA were amplified by PCR using primers that would bind to a CpG-free sequence from the MYOD1 gene only after complete bisulfite conversion of all cytosines, thus allowing assessment of the efficacy of bisulfite conversion. *MYOD1* primer set was 5’-CCAACTCCAAATCCCCTCTCTAT-3’ (forward), and 5’-TGATTAATTTAGATTGGGTTTAGAGAAGGA-3’ (reverse). 9 μl of the bisulfite converted DNA was subjected to nested PCR for *GSTP1* alleles. The outer PCR was carried out in 40 μl reactions containing 10x Platinum Taq buffer (Invitrogen, Carlsbad, CA), 1.5 U Platinum Taq (Invitrogen), 250 μM each dNTPs, 1.5 mM MgCl2, 0.25 μg/μl BSA, 2 μl dimethyl sulfoxide, 400 nM forward primer and 400 nM reverse primer. Cycling conditions were 95°C for 3 min, 40 cycles of 95°C for 30 s, 50°C for 45 s and 72°C for 45 s, followed by a 7 min extension step at 72°C. The outer PCR products amplified using primer #1 and #2 were purified and concentrated using the DNA Clean and Concentrator 5 kit (Zymo Research, Orange, CA) and eluted in 20ul of TE buffer. Aliquots (4 μl) of the cleaned PCR product used in the inner PCR amplification using the #3 and #4 primers in a 40 μl reaction mixture for 20 cycles. Primers amplifying *GSTP1* CpG islands without bias to methylation patterns were #1 forward primer, 5’-GTTGGTTTTATGTTGGGAGTTTTGAG-3’; #2 reverse primer, 5’-ATCCTCTTCCTACTATCTATTTACTCCCTAA-3’; #3 forward primer, 5’-GTTGGGAGTTTTGAGTTTTATTTT-3’; and #4 reverse primer, 5’-ACTATCTATTTACTCCCTAAACCCC-3’. Nested PCR products were gel purified and cloned into pCR^®^2.1-TOPO^®^ vectors (Invitrogen, Carlsbad, CA). Plasmid DNA from 10 positive clones was sequenced using an ABI 3730 DNA Analyzer (Applied Biosystems, Foster City, CA). Bisulfite sequencing data were analyzed using a custom Java program called DNAMethylMap, which facilitates the analysis of Sanger bisulfite sequencing clones with virtually bisulfite converted reference amplicon sequences (17). Clones that showed conversion of >95% of cytosines outside of a CpG context were considered valid. DNA from the prostate cancer cell line LNCaP (RRID:CVCL_0395), which is known to be completely methylated at all CpGs in the region analyzed (18), was used as a positive control for methylation, and normal human WBC DNA (Novagen, Madison, WI) was used as a negative control.

### Statistical Assessment

The overall extent of *GSTP1* gene promoter methylation across the four tissue types (normal, PIA, HGPIN, and CaP) were assessed with the Mann-Whitney U test. The extent of *GSTP1* CpG dinucleotide methylation on an individual clone/allele across all 39 CpGs analyzed for that allele was classified as negative (<10%), mild (10-25%), moderate (25-50%) and high (>50%).

Differences in frequency of *GSTP1* hypermethylation between normal, atrophy, HGPIN, and CaP were assessed with the χ^2^ test, Fisher’s exact test, and Mann-Whitney U test. For further interactive analysis and visualization of the data, we used a custom Java program called DNAMethylMap v1.2 (17).

## Results

### Laser capture microdissection and bisulfite sequencing of the GSTP1 promoter CpG island

The *GSTP1* gene is approximately 4 kb in length, comprises 7 exons and 6 introns (19,20) (Fig. 1A). Primers amplifying the *GSTP1* CpG island without bias to methylation patterns allowed investigation into the methylation status of 39 CpG dinucleotides between −254 to +81 upstream of the *GSTP1* promoter region (GenBank accession number AY324387, [GRCh38, Chr11:67,583,561-67,583,896]) (Fig. 1A).

**Figure 1.**
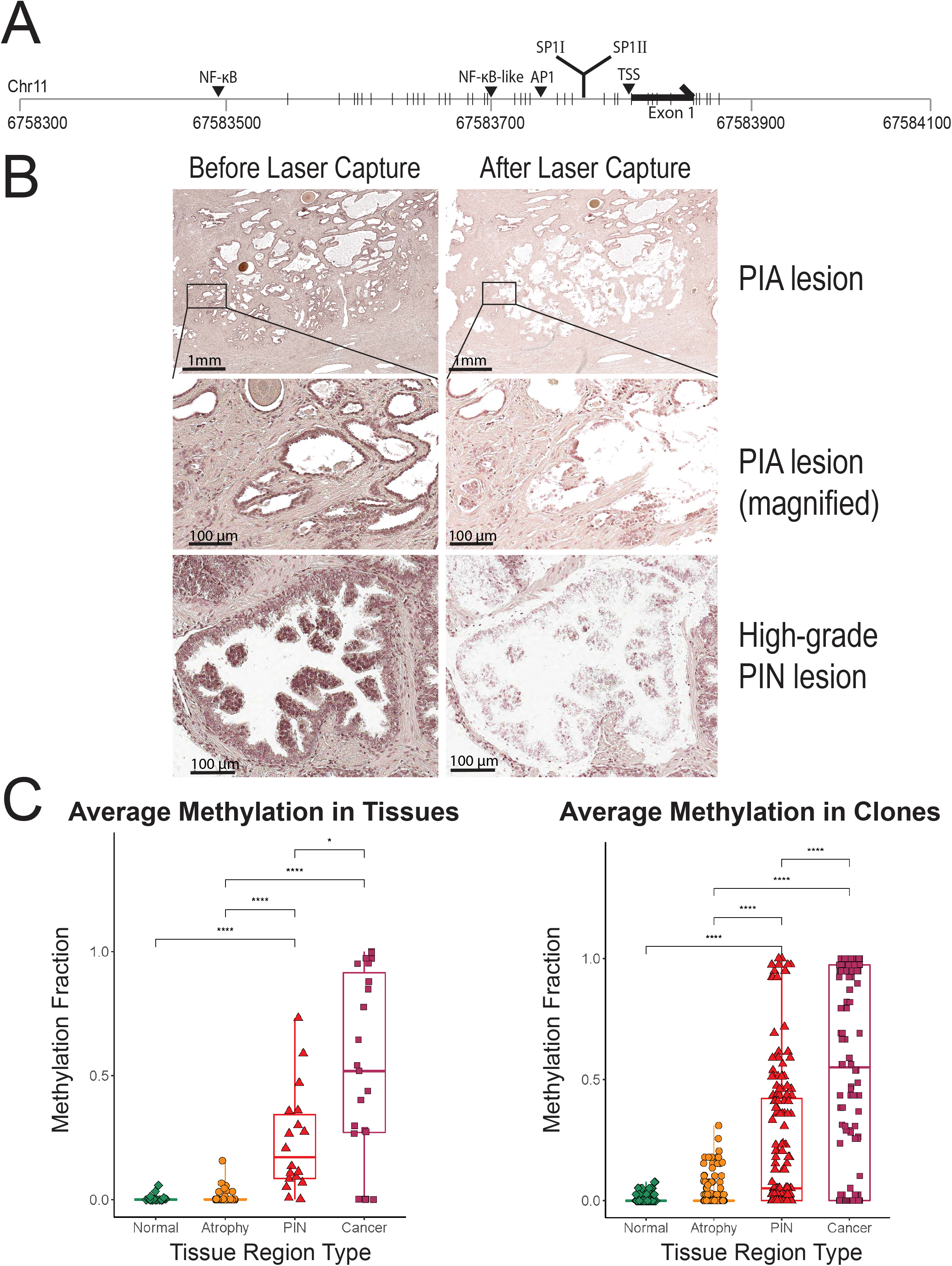
Characterization of GSTP1 promoter methylation using bisulfite sequencing of 39 CpGs within and surrounding the core GSTP1 promoter in micro-dissected tissues. (A) Schematic of the GSTP1 promoter locus with positions of the interrogated CpGs (marked with vertical black line) and notable landmark positions (NF-kB binding site, AP1 binding site, SP1I binding site, SP1II binding site, and transcription start site). (B) H&E view of a representative case showing a region of prostatic inflammatory atrophy (PIA) and a high-grade PIN lesion before and after laser capture microdissection. In the middle panels of PIA, the arrows indicate region of transition to cells with low level nuclear atypia, and the arrow heads indicate an atypical gland likely budding from the same lesion, which was purposely not procured during laser capture microdissection. (C) Methylation of the GSTP1 promoter averaged across all the clones in each of 102 lesions (left), and percent methylation of the GSTP1 promoter for each of 908 individual cloned amplicons (right). Mann-Whitney U test was used to assess statistically significant differences in plots. Asterisks denote level of significance (* p < 0.05; ** p < 0.01; *** p < 0.001; **** p < 0.0001).

To obtain genomic DNA from prostate tissues enriched for epithelial cells from normal prostate, proliferative inflammatory atrophy lesions (PIA), high grade prostatic intraepithelial neoplasia (HGPIN), and prostate adenocarcinoma (CaP), laser capture microdissection was undertaken using radical prostatectomy specimens from 102 lesions across 32 patients; representative images of PIA and PIN lesions before and after LCM are shown in Fig. 1B. Using nested amplification of bisulfite-treated DNA followed by cloning of individual PCR products, sequencing for ^5-me^C versus C was performed on individual clones/alleles from *GSTP1* promoter amplicons. A total of 212 clones from normal epithelium (24 LCM regions), 327 clones from PIA (37 LCM regions), 167 clones from HGPIN (18 LCM regions), and 202 clones from CaP (23 LCM regions) were sequenced, giving a sum total of 34,863 CpG sites analyzed. The promoter region of *GSTP1* in the LNCaP human CaP cell line was almost completely methylated (5644/5770 or 98%), while white blood cells showed virtually no CpG methylation (24/5670 or 0.42%), as expected.

### GSTP1 promoter CpG island methylation patterns in normal glands, PIA, PIN, and invasive cancer lesions

Overall, a gradual increase in the average methylation level was evident across normal, PIA, PIN, and cancer regions (Fig. 1C). The difference in average percent methylation of the *GSTP1* promoter was significant between normal prostate and HGPIN (p < 0.0001, Mann Whitney U test), and between PIA and HGPIN (p < 0.0001, Mann Whitney U test, Fig. 1C).

We next examined the patterns of methylation at each individual allele in each tissue type (Fig. 2). Normal prostate epithelium was almost completely unmethylated (42/8079 CpGs methylated or 0.52%, Fig. 2A), and all clones had less than 10% methylation (Fig. 3). In PIA lesions, very sparse *GSTP1* promoter methylation was evident (164/12521 or 1.31%; Fig. 2B) with only 7 of 31 lesions, representing 18 of 327 clones, showing any evidence of methylation (Supplementary Table 3). One of the PIA lesions showed one clone with moderate methylation. Additionally, 6 of 7 PIA lesions with >10% methylated CpGs were either next to OR merging with PIN or atypical glands suspicious for, but not diagnostic of, cancer (e.g. often referred to as “atyp” or atypical small acinar proliferation [ASAP]), compared to 10 of 30 PIA lesions with <10% methylated CpGs (p = 0.0118, χ^2^ test). The pattern of methylation of *GSTP1* alleles in PIA lesions showed only sparse methylation, usually having less dense methylation changes compared to PIN and CaP lesions (Fig. 2).

**Figure 2.**
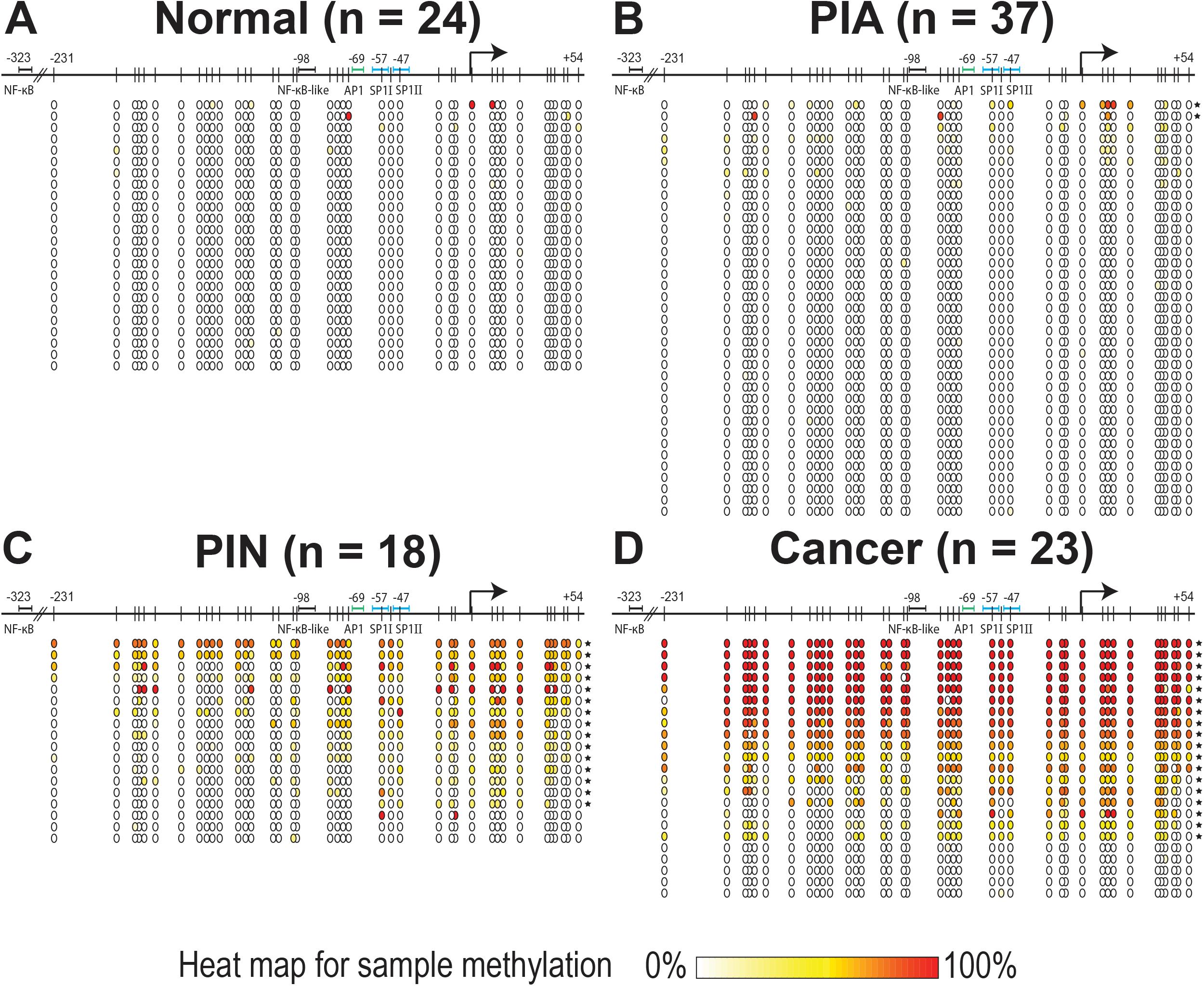
CpG methylation heat maps of average methylation across all alleles in a lesion for each CpG in various prostatic tissues. Each circle corresponds to a CpG, and the color indicates the percent of clones methylated at that CpG in the given lesion. Each row represents one region isolated by laser capture microdissection and subjected to bisulfite sequencing. Starred rows indicate a lesion where the methylation average across all clones and all CpGs for the given lesion is greater than 6%. (A) Normal prostatic epithelium, (B) prostatic inflammatory atrophy (PIA). (C) Prostatic intraepithelial neoplasia (PIN), and (D) prostatic adenocarcinoma.

**Figure 3.**
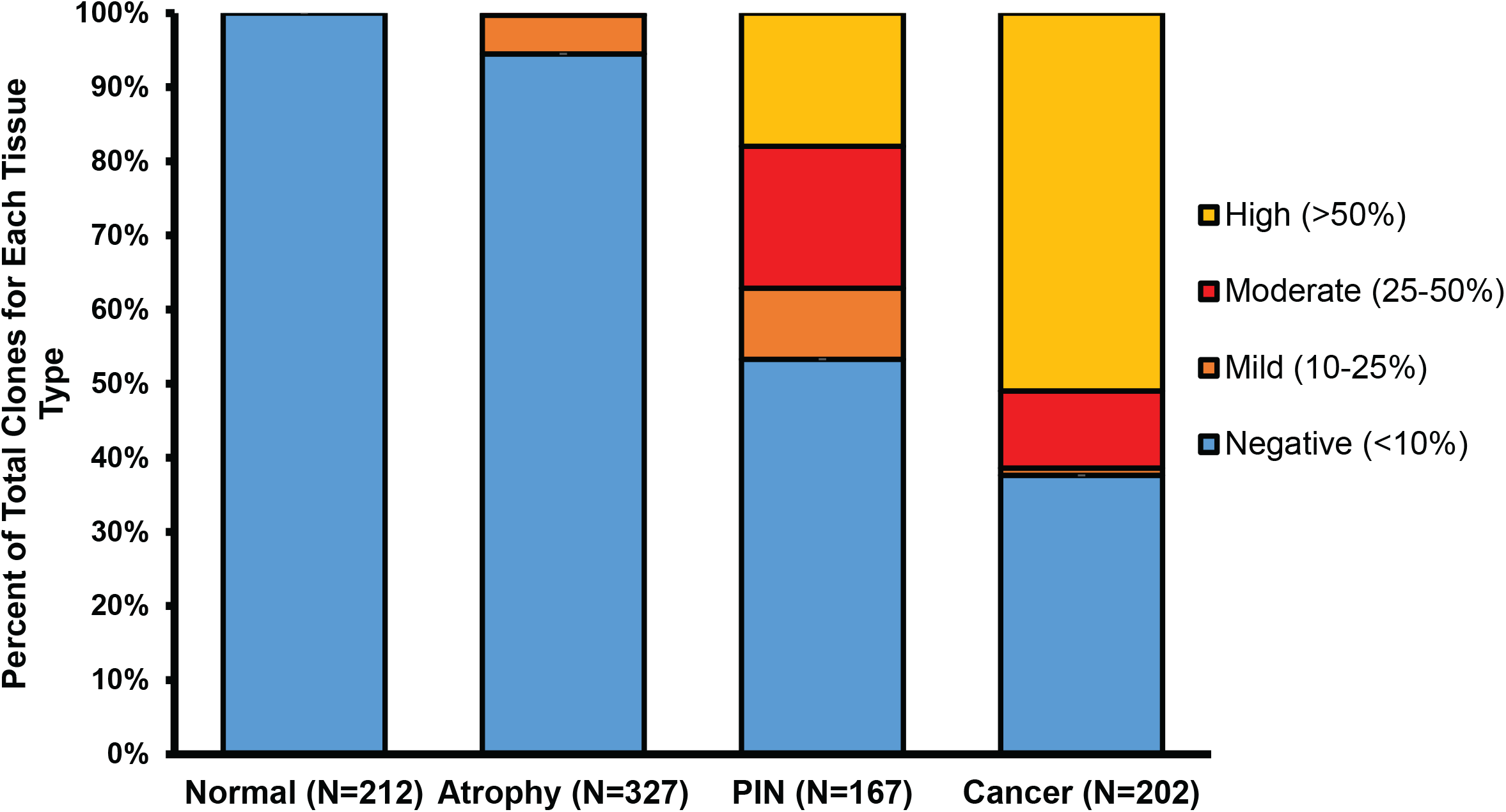
Phased methylation by allele of the GSTP1 promoter region. For each of the 908 alleles assessed across 102 lesions, the extent of *GSTP1* CpG dinucleotide methylation on an individual allele was classified as negative, mild, moderate, or high if the proportion of methylated CpGs in the allele represented <10%, 10-25%, 25-50%, or >50% of the total CpGs in that allele, respectively.

HGPIN lesions showed an intermediate level of *GSTP1* promoter methylation between normal/PIA and CaP. HGPIN had 1534 of 6461 (24%) CpGs methylated (Fig. 2C). 28.7% of PIN alleles had a mild or moderate level of methylation that was significantly different from the 5.5% in PIA (p = 6.9E-13, χ^2^ test) and 11.4% in cancer (p = 2.6E-05, χ^2^ test). Many individual alleles in HGPIN showed an intermediate density of methylation compared to normal/PIA and to CaP (Fig. 2C). CaP lesions showed an increase in CpG methylation level and density over that in HGPIN, with 4002 of 7802 (51.3%) CpG sites showing methylation and 124 of 202 individual *GSTP1* alleles (61.4%) exhibiting moderate and high density of methylation. The increase in highly methylated *GSTP1* promoter alleles in CaP vs HGPIN was significant (p = 3.5E-06, χ^2^ test).

*GSTP1* promoter CpG sites were more commonly methylated near the transcription start site and transcription factor binding sites for AP1 and SP1 than in further upstream sequences (Fig. 2). Six proximally located CpG sites (between −4 and +38 of the transcription start site) could be considered “hot spots” for somatic methylation changes, appearing methylated in *GSTP1* with either sparse, intermediate, or dense methylation patterns from atrophy, HGPIN, and CaP (Fig. 2). There was also a smaller upstream region (−197 to −176) with recurrently methylated CpGs not correlating with any known transcription factor binding sites.

### Allele level intra- and inter-patient heterogeneity in GSTP1 promoter CpG island methylation

To highlight the patterns of intra- and inter-individual heterogeneity, we carefully examined four cases in which genomic DNA was isolated from laser capture microdissected normal, PIA, PIN, and invasive cancer regions (Fig. 4–6). In all four of these patients, we observed very minimal methylation in all individual alleles in both the normal and PIA lesions. In comparing patients 21964 and 1618, both cases had highly dense methylation across all the *GSTP1* alleles in CaP and showed intermediate *GSTP1* methylation in the PIN lesions (Fig. 4). However, the PIN lesion in patient 21964 had alleles with heterogeneity in the *GSTP1* promoter methylation patterns (Fig. 4A), but patient 1618 had nearly identical methylation across all 10 clones from the PIN lesion (Fig. 4B).

**Figure 4.**
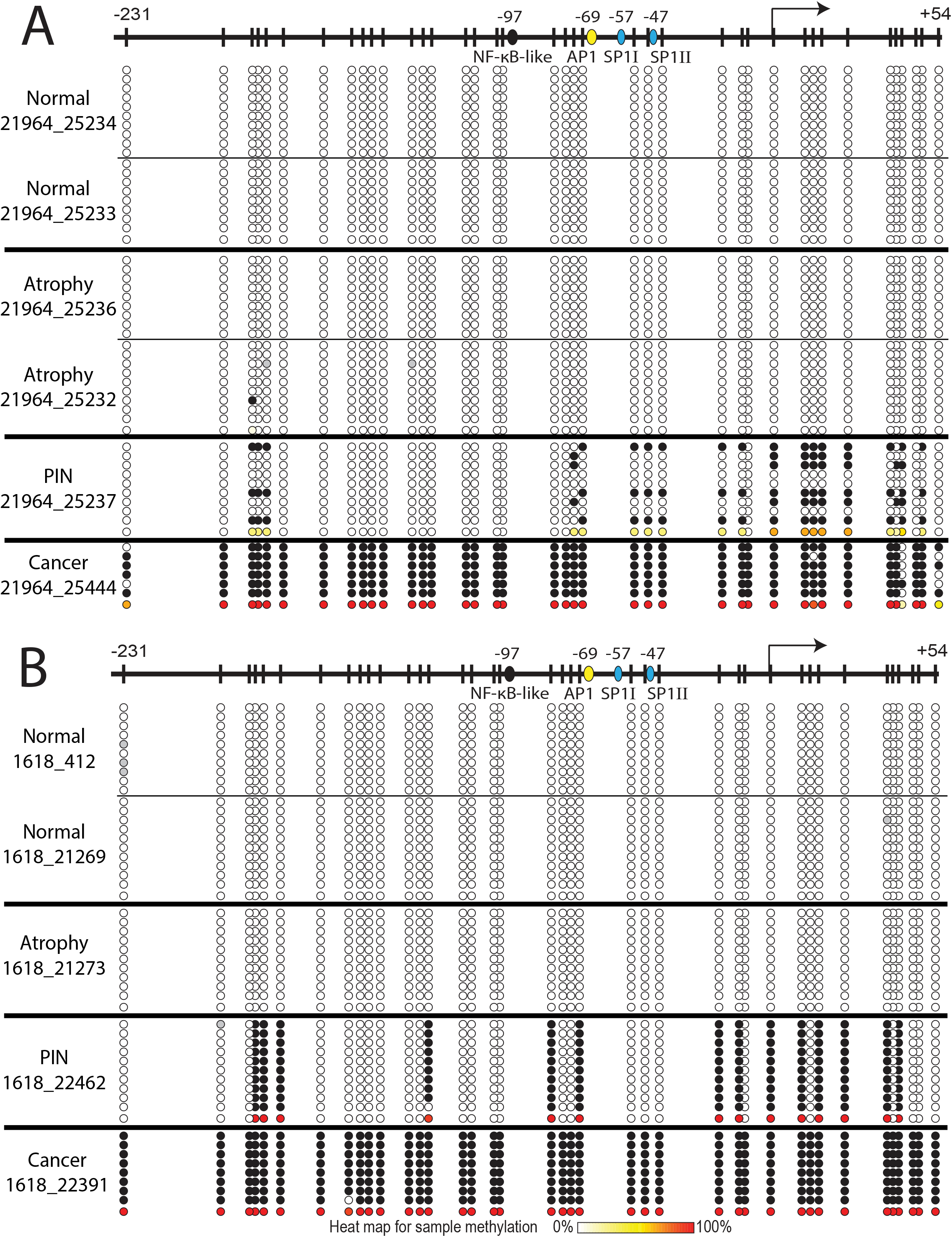
Intra-patient and inter-patient heterogeneity in allele-level GSTP1 promoter methylation patterns for patient cases 21964 and 1618 with all lesion types microdissected. For every lesion that was laser capture microdissected from each patient, each allele is represented on a given line with the methylation status of the 39 CpGs in the GSTP1 promoter. For each allele, a black circle represents a methylated CpG, a white circle an unmethylated CpG, and a gray circle is unknown. The bottom row in each lesion subtype (normal, atrophy, PIN, cancer) represents the methylation for each CpG averaged across all the alleles for the lesion. (A) Patient 21964, (B) Patient 1618

In examining the *GSTP1* alleles for CaP lesions in Patient 28126, an averaged methylation view signifies an intermediate level of methylation (Fig. 5). However, there were clearly two distinct groups of alleles, 5 clones with nearly complete methylation and the other four without any methylation (Fig. 5). Unlike the previous two patients (Fig. 4), the PIN lesion from this subject had a similar methylation pattern to the CaP lesion (Fig. 5). This pattern of having two distinct fully methylated or unmethylated allele subsets was observed in a total of 3 of 23 CaP lesions.

**Figure 5.**
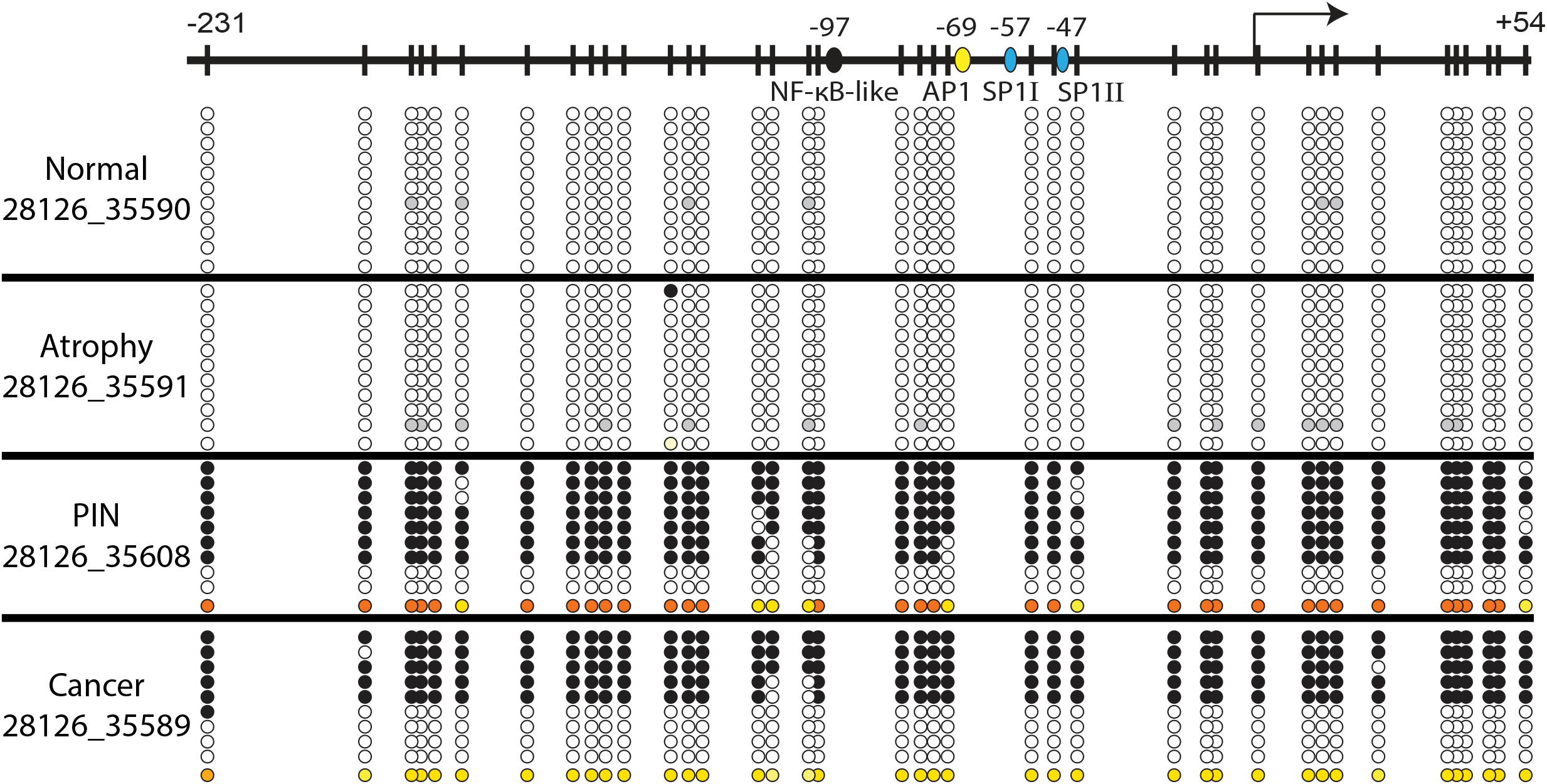
Patient 28126 allele-level GSTP1 promoter methylation patterns. For every lesion that was laser capture microdissected from each patient, each allele is represented on a given line with the methylation status of the 39 CpGs in the GSTP1 promoter. For each allele, a black circle represents a methylated CpG, a white circle an unmethylated CpG, and a gray circle is unknown. The bottom row in each lesion subtype (normal, atrophy, PIN, cancer) represents the methylation for each CpG averaged across all the alleles for the lesion.

In patient 26559, it is notable that three different cancer lesions harbored three distinct methylation patterns (Fig. 6). One cancer lesion had clones with full methylation across the *GSTP1* promoter, the next one had clones with intermediate levels of methylation, and the last cancer lesion had no methylation across all of the clones. This highlights the degree of heterogeneity there can be with the *GSTP1* CpG island promoter hypermethylation patterns within different tumor sites in a given patient.

**Figure 6.**
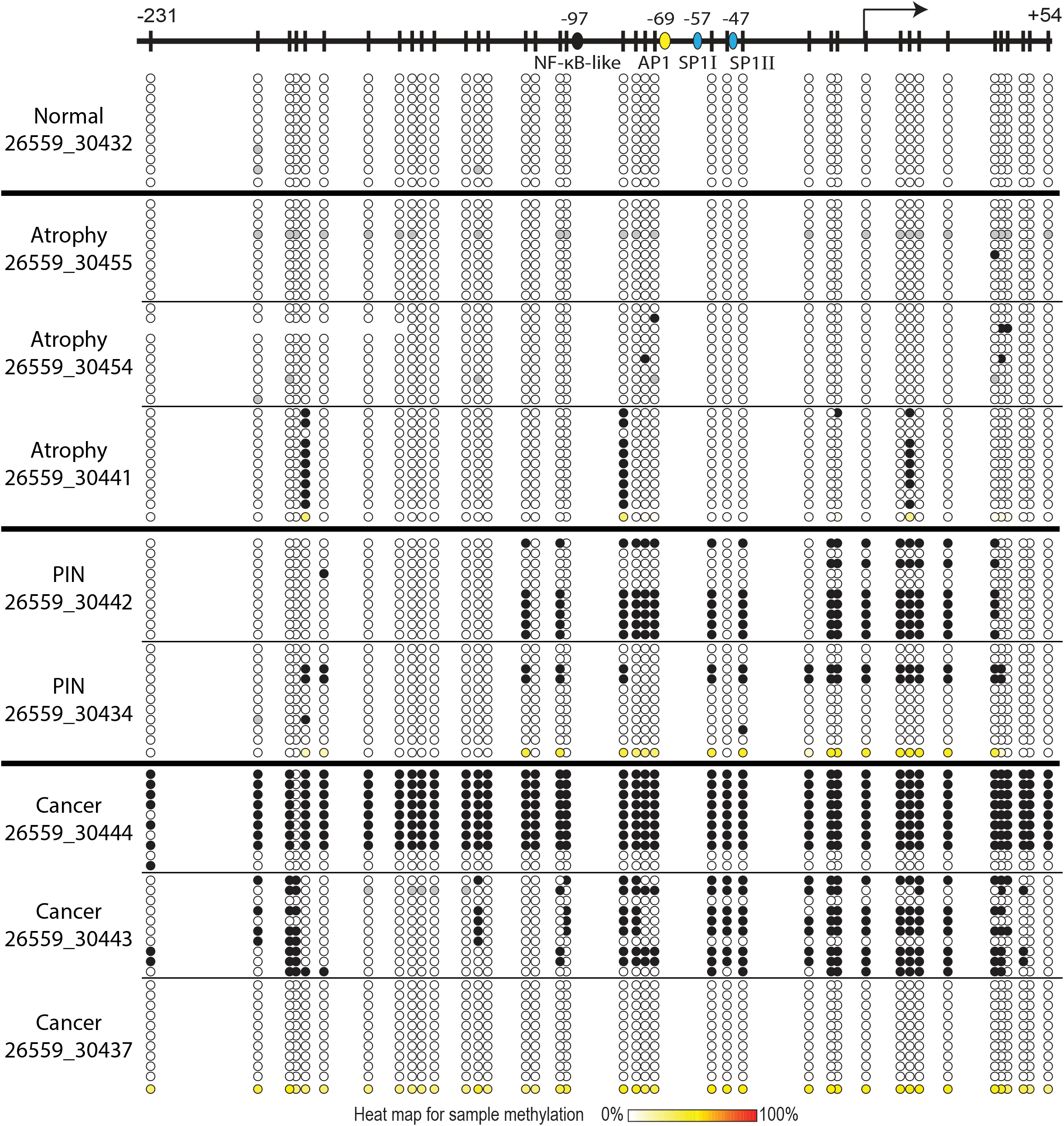
Patient 26559 allele-level GSTP1 promoter methylation patterns. For every lesion that was laser capture microdissected from each patient, each allele is represented on a given line with the methylation status of the 39 CpGs in the GSTP1 promoter. For each allele, a black circle represents a methylated CpG, a white circle an unmethylated CpG, and a gray circle is unknown. The bottom row in each lesion subtype (normal, atrophy, PIN, cancer) represents the methylation for each CpG averaged across all the alleles for the lesion.

The expression of GSTP1 protein as detected by immunohistochemistry in Patient 21964 demonstrates that lack of methylation in the *GSTP1* promoter (Fig. 4A) corresponds with GSTP1 expression in normal and PIA lesions (Fig. 7A-D). Additionally, the presence of hypermethylation (Fig. 4A) corresponds with silencing of GSTP1 in luminal cells with GSTP1 expression present only in stromal cells in the cancer lesion (Fig. 7G-H). The PIN allele-level distribution shows 3 alleles without and 6 clones with methylation (Fig. 4A), likely corresponding with the distinct regions of GSTP1-positive and GSTP1-negative expression respectively (Fig. 7E-F). In a different patient, a similar pattern of GSTP1 expression heterogeneity corresponding with *GSTP1* methylation heterogeneity is observed in an HGPIN lesion and in a PIA lesion, in which a few regions of low levels of GSTP1 expression are demonstrated (Supplementary Fig. 1).

**Figure 7.**
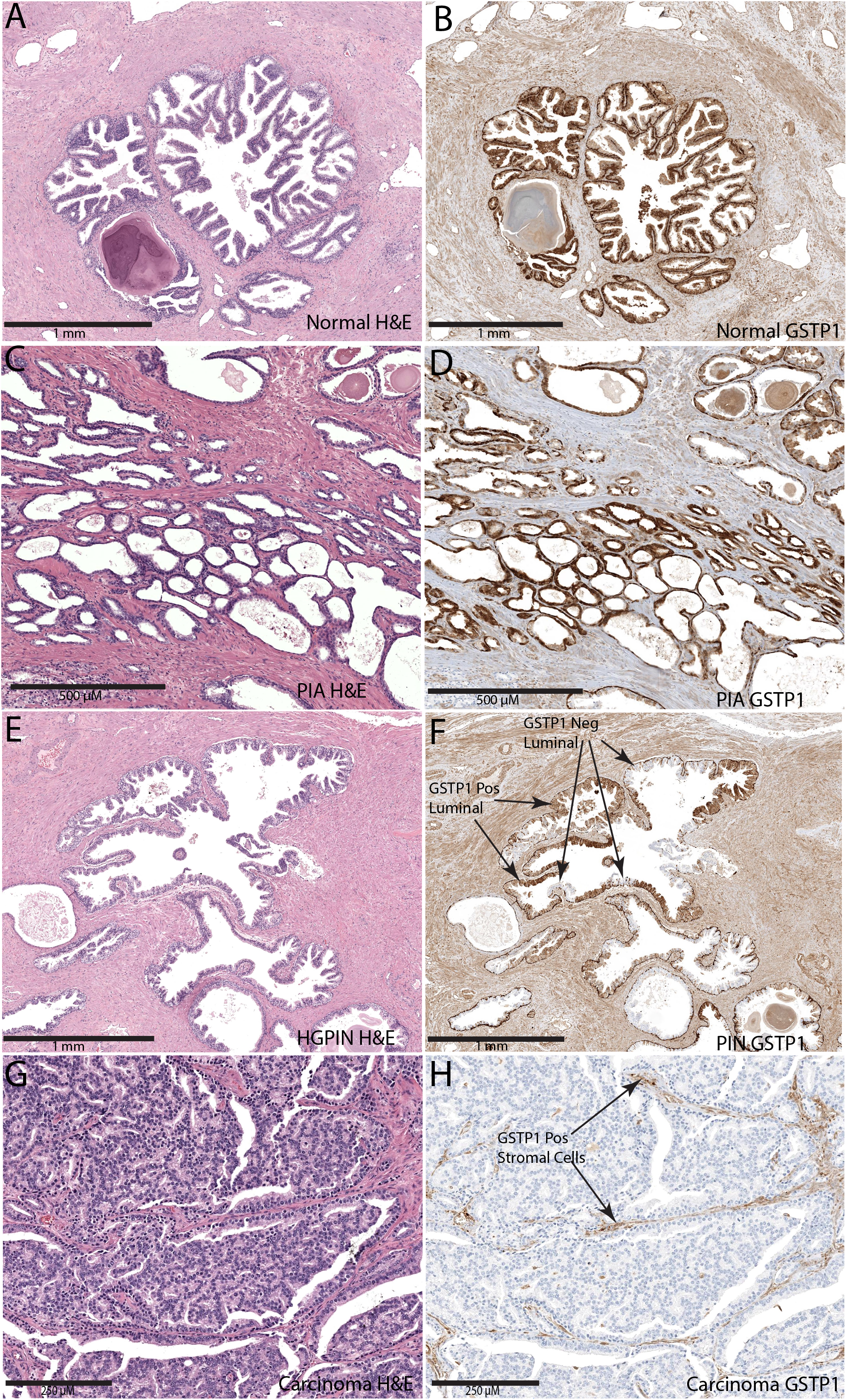
GSTP1 expression assessed by immunohistochemistry (IHC) on laser capture microdissected prostate glands in each tissue type for patient 21964. (A) H&E of normal prostatic epithelium, (B) GSTP1 IHC of normal prostatic epithelium,(C) H&E of prostatic inflammatory atrophy (PIA), (D) GSTP1 IHC of PIA, (E) H&E of high-grade prostatic intraepithelial neoplasia (HGPIN), (F) GSTP1 IHC of HGPIN, (G) H&E of prostatic adenocarcinoma, and (H) GSTP1 IHC of prostatic adenocarcinoma.

## Discussion

Somatic CpG island hypermethylation, which results in gene silencing, accompanies human prostate cancer pathogenesis, and is often evident in early stages of disease development, including cancer precursor lesions (4). Intraepithelial neoplasia (IEN) refers to epithelial lesions in many organ sites that exhibit cytological features of carcinoma but fail to invade through basement membrane structures and are thought to represent precursors to cancer, and have been proposed to represent conditions suitable for therapeutic intervention in pursuit of cancer prevention or interception (21). In the prostate, HGPIN is an IEN lesion that is traditionally thought to serve as a precursor to many CaP lesions. In this study, the levels and density of *GSTP1* CpG island hypermethylation in HGPIN lesions were often intermediate between normal prostate epithelium and CaP. This observation provides molecular evidence that HGPIN lesions often represent an intermediate step in the pathogenesis of CaP. Interestingly, the levels of *GSTP1* CpG island hypermethylation in PIA lesions showed a trend to be higher than normal, but significantly less dense than HGPIN. However, since CaP cells can grow along established prostate glands and duct structures, some fraction of HGPIN lesions could in fact represent CaP mimicking IEN (11,12). The identical *GSTP1* promoter CpG island methylation patterns between CaP and HGPIN lesions in the same patient (Fig. 5) may represent such cases of retrograde spread of invasive carcinomas into prostate glands. Conversely, this could also represent a case in which the PIN had “progressed” in methylation and gave rise to this invasive carcinoma.

Methylation-specific primers (MSP), and related techniques, are used to detect methylation of CpG dinucleotides at specific genome sites. MSP assays have reported a limit of detection below 0.1% methylated alleles (22). Using MSP, CpG methylation levels are directly assessed only at primer annealing sites, so using different MSP primer sets, the frequency of *GSTP1* promoter methylation levels in different prostate lesions from different prostate cancer cases has appeared variable throughout different studies (15). The different primer sets are prone to false negative results depending on the distribution of methylated CpGs across the *GSTP1* promoter (4,15,18). The bisulfite genomic sequencing assay strategy used in this study was capable of providing the pattern and phasing of 5-meCpG along individual alleles. In this strategy, bisulfite treatment of genomic DNA is combined with the sequencing of individual product clones generated by PCR amplification (23). The method also permits monitoring of conversion efficiency for unmethylated cytosine bases; PCR product clones from a bisulfite genomic sequencing assay can be discarded if inadequately converted. The major limitation of the bisulfite genomic sequencing approach appears to be DNA fragmentation during the bisulfite reaction, with as much as 90 ± 6% of input DNA subject to some level of degradation during bisulfite treatment (24). Hence, although it is possible to amplify sequences from as few as 10 copies of bisulfite-treated genomes in theory, the consistency and quality of the bisulfite genomic sequencing results tend to be markedly better if more DNA is used. For our study, we collected as much as 2 ng of genomic DNA and were successfully able to recover bisulfite converted *GSTP1* promoter sequences as PCR clones. However, even though we sequenced 10 clones per sample for the study, the PCR products could have been derived from as few as 1 or 2 DNA molecules, resulting in stochastic clonal amplification (25). Thus, observing identical methylation patterns across clones (for example, the PIN lesion from patient 28126, Fig. 4B) may either indicate a true homogeneity in the methylation pattern in that sample, or could be due to clonal amplification of a single allele. Despite this limitation, the insights generated by examining the density of CpG methylation on individual *GSTP1* alleles were very informative: among all *GSTP1* alleles sequenced, the density of methylation in genomic DNA from HGPIN was clearly intermediate between normal prostate/PIA and CaP.

Among *GSTP1* alleles from prostate cancer precursor lesions subjected to bisulfite genomic sequencing, an apparent “hot spot” for CpG methylation surrounding the transcription start site and the AP1/SP1 binding sites was identified. In an analysis of de novo CpG methylation in transfected *GSTP1* promoters in prostate cancer cells, Song et al. observed a spread of cytosine methylation with serial passage of the cells if (i) an initial 5-meCpG was present to “seed” the allele, and (ii) transcription from the allele was silenced (26). In their experiments, CpGs immediately downstream of the transcription start site (and near the “hot spot” seen in the current study) most commonly showed *de novo* methylation. A seed and spread model may be able to account for the apparent increase in the CpG methylation density in *GSTP1* alleles during prostatic carcinogenesis. Rare *GSTP1* CpG dinucleotides might become methylated spontaneously or in response to stress in normal prostate epithelial cells/atrophy cells, and then additional methylated CpG dinucleotides might accumulate in the setting of transcriptional silencing. Perhaps, CpG methylation at the “hot spot” itself contributes to inhibition of transcription, serving to drive both seeding and spreading of a wave of cytosine methylation. The intrapatient heterogeneity of *GSTP1* promoter CpG island methylation of CaP lesions (Fig. 6), may represent three events in the spectrum of such a seed and spread model. The observed cancer lesion devoid of *GSTP1* promoter CpG methylation could represent an early cancer event in which no seeding event had yet occurred, while the cancer with intermediate levels of methylation contains cells soon after initial seeding and spread of CpG methylation around the “hotspot” area (Fig. 6). The cancer lesion with complete *GSTP1* promoter methylation may indicate complete progression once the CpG methylation has spread throughout the entire *GSTP1* promoter CpG island. The findings from this study raise new questions as to whether other densely methylated genes in prostate cancer show similar “seeding and spreading” patterns in prostate cancer precursor lesions.

## Acknowledgements

We thank the Christina DeStefano Shields and Brenna Hairston for critical reading of the manuscript.

