## Supplementary Figures and Tables for "Progressive spreading of DNA methylation in the GSTP1 promoter CpG island across transitions from precursors to invasive prostate cancer"

### Supplementary Materials

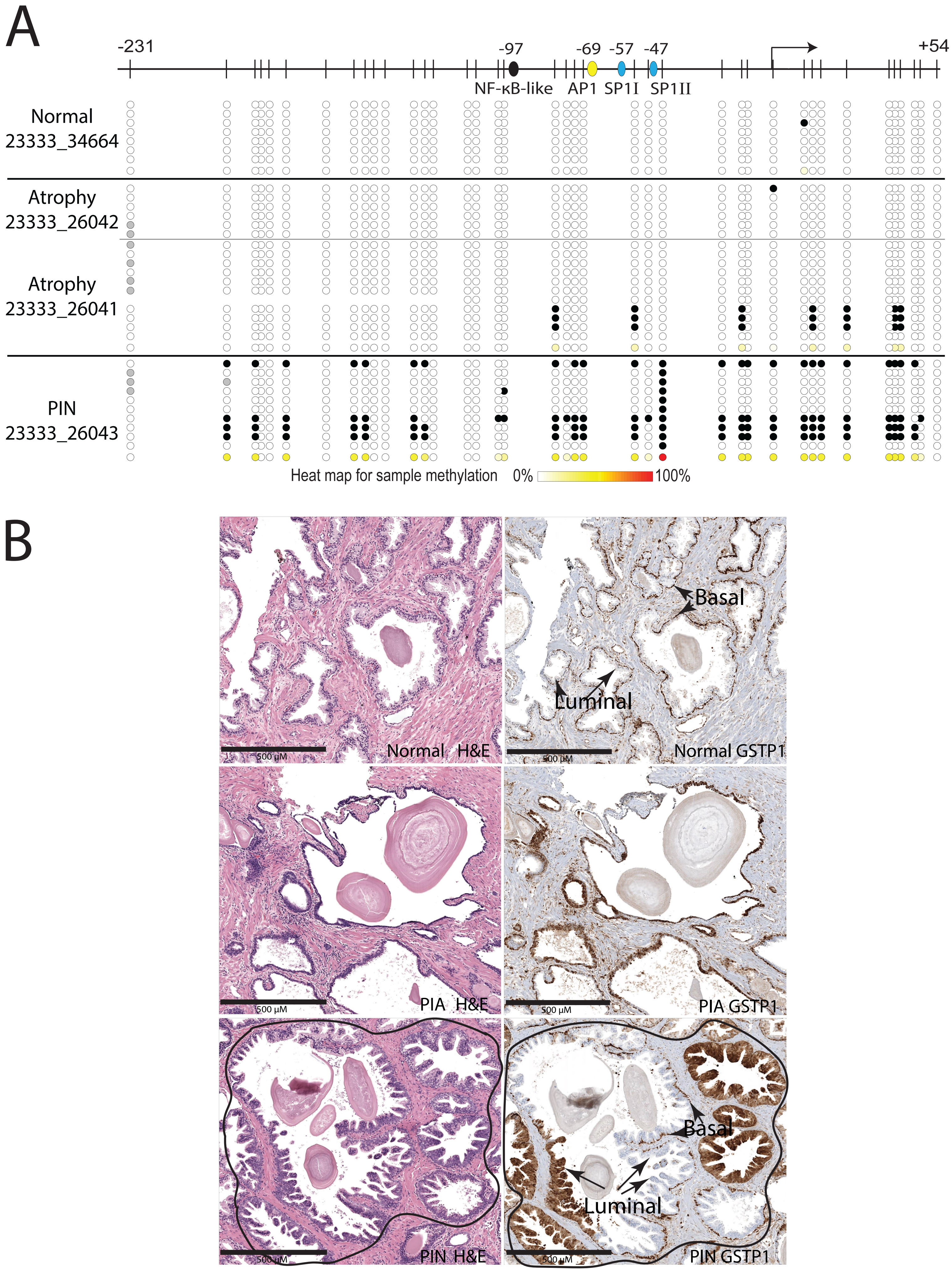
A
-231
-97
-69
-57
-47
+54
NF-κB-like
AP1
SP1I
SP1II
Normal
23333_34664
Atrophy
23333_26042
Atrophy
23333_26041
PIN
23333_26043
Heat map for sample methylation
0%
100%
B
Basal
Luminal
500 μM
Normal H&E
Normal GSTP1
PIA H&E
PIA GSTP1
PIN H&E
PIN GSTP1

**Supplementary Figure 1**. **Intra-patient heterogeneity in GSTP1 methylation and GSTP1 expression in patient 23333.** (A) For every lesion that was laser capture microdissected from this patient, each allele is represented on a given line with the methylation status of the 39 CpGs in the GSTP1 promoter. For each allele, a black circle represents a methylated CpG, a white circle an unmethylated CpG, and a gray circle is unknown. The bottom row in each lesion subtype (normal, atrophy, PIN) represents the methylation for each CpG averaged across all the alleles for the lesion. (B). Corresponding H&E and immunohistochemistry for GSTP1.

| **Characteristic** | **N = 32**[1] |
|---|---|
| **Age at Prostatectomy (years)** | 59.4 (48.0,69.0) |
| **Race** | |
| Black or African American | 1 (3.1%) |
| White or Caucasian | 31 (97%) |
| **Surgical Margin** | |
| Negative | 30 (94%) |
| Positive | 2 (6.2%) |
| **Mean Weight of Prostate Gland (grams)** | 54 (36,95) |
| **Gleason Grade Group** | |
| Grade Group 1 | 23 (74%) |
| Grade Group 2 | 1 (3.2%) |
| Grade Group 3 | 4 (13%) |
| Grade Group 4 | 1 (3.2%) |
| Grade Group 5 | 2 (6.5%) |
| N/A | 1 |
| **Tumor Stage** | |
| T0N0MX | 1 (3.1%) |
| T2N0MX | 24 (75%) |
| T3AN0MX | 3 (9.4%) |
| T3AN1MX | 1 (3.1%) |
| T3BN0MX | 3 (9.4%) |

[1] Mean (Minimum,Maximum); n (%)

**Supplementary Table 1. Patient and tumor characteristics of 32 patients who underwent radical prostatectomies.**

| Observed Frequency | Normal | Atrophy | PIN | Cancer | **Total** |
|---|---|---|---|---|---|
| Negative (<10%) | 24 | 30 | 3 | 5 | 62 |
| Mild (10-25%) | 0 | 6 | 2 | 0 | 8 |
| Moderate (25-50%) | 0 | 1 | 7 | 2 | 10 |
| High (>50%) | 0 | 0 | 6 | 16 | 22 |
| **Total** | 24 | 37 | 18 | 23 | 102 |

**Supplementary Table 2. Phased methylation of the GSTP1 promoter region across 102 lesions.** For each of the 102 lesions assessed across the four tissue types, the extent of *GSTP1* CpG dinucleotide methylation in a given lesion was classified as negative, mild, moderate, or high if the proportion of methylated CpGs across all clones in that lesion represented <10%, 10-25%, 25-50%, or >50% of the total CpGs, respectively.

| Observed Frequency | Normal | Atrophy | PIN | Cancer | **Total** |
|---|---|---|---|---|---|
| Negative (<10%) | 212 | 309 | 89 | 76 | 686 |
| Mild (10-25%) | 0 | 17 | 16 | 2 | 35 |
| Moderate (25-50%) | 0 | 1 | 32 | 21 | 54 |
| High (>50%) | 0 | 0 | 30 | 103 | 133 |
| **Total** | 212 | 327 | 167 | 202 | 908 |

**Supplementary Table 3. Phased methylation of the GSTP1 promoter region across 908 clones.** For each of the 908 alleles assessed across the four tissue types, the extent of *GSTP1* CpG dinucleotide methylation on an individual allele was classified as negative, mild, moderate, or high if the proportion of methylated CpGs in the allele represented <10%, 10-25%, 25-50%, or >50% of the total CpGs in that allele, respectively.
